# Glioblastoma Mesenchymal Transition and Invasion are Dependent on a NF-κB/BRD2 Chromatin Complex

**DOI:** 10.1101/2023.07.03.546613

**Authors:** Raghavendra Vadla, Shunichiro Miki, Brett Taylor, Daisuke Kawauchi, Brandon M Jones, Nidhi Nathwani, Philip Pham, Jonathan Tsang, David A. Nathanson, Frank B. Furnari

## Abstract

Glioblastoma (GBM) represents the most aggressive subtype of glioma, noted for its profound invasiveness and molecular heterogeneity. The mesenchymal (MES) transcriptomic subtype is frequently associated with therapy resistance, rapid recurrence, and increased tumor-associated macrophages. Notably, activation of the NF-κB pathway and alterations in the *PTEN* gene are both associated with this malignant transition. Although PTEN aberrations have been shown to be associated with enhanced NF-κB signaling, the relationships between PTEN, NF-κB and MES transition are poorly understood in GBM. Here, we show that PTEN regulates the chromatin binding of bromodomain and extraterminal (BET) family proteins, BRD2 and BRD4, mediated by p65/RelA localization to the chromatin. By utilizing patient-derived glioblastoma stem cells and CRISPR gene editing of the *RELA* gene, we demonstrate a crucial role for RelA lysine 310 acetylation in recruiting BET proteins to chromatin for MES gene expression and GBM cell invasion upon *PTEN* loss. Remarkably, we found that BRD2 is dependent on chromatin associated acetylated RelA for its recruitment to MES gene promoters and their expression. Furthermore, loss of BRD2 results in the loss of MES signature, accompanied by an enrichment of proneural signature and enhanced therapy responsiveness. Finally, we demonstrate that disrupting the NF-κB/BRD2 interaction with a brain penetrant BET-BD2 inhibitor reduces mesenchymal gene expression, GBM invasion, and therapy resistance in GBM models. This study uncovers the role of hitherto unexplored PTEN-NF-κB-BRD2 pathway in promoting MES transition and suggests inhibiting this complex with BET-BD2 specific inhibitors as a therapeutic approach to target the MES phenotype in GBM.

## Introduction

Glioblastoma (GBM), the most aggressive and malignant form of glioma, is particularly notorious for its ability to invade healthy brain tissue beyond the visible tumor margins (1). This invasion is mediated by a complex interplay of molecular signaling pathways and cellular interactions between tumor cells and the surrounding microenvironment (2). A central issue confounding successful treatment is the heterogeneous nature of this aggressive tumor. Multi-omics analyses identified three transcriptomic subtypes in IDHwt GBM, classical (CL), mesenchymal (MES), and proneural (PN), with individual tumors typically harboring mixtures of all three subtypes (3,4). The mesenchymal subtype is associated with therapy resistance, increased invasion, and presence of tumor associated macrophages (TAMs) (5). Recent studies have shown that IDHwt tumors with a mesenchymal recurrence exhibited a significantly shorter surgical interval compared with those with non-mesenchymal recurrence(6). These characteristics highlight a need to better understand the factors that drive mesenchymal transition in GBM.

Activation of the NF-κB pathway plays a crucial role in promoting MES transition by upregulating the expression of several key mesenchymal genes and also by stimulating the expression of various cytokines, chemokines, and growth factors, thereby further contributing to the mesenchymal phenotype in GBM cells (7,8). PTEN, an important negative regulator of the NF-κB pathway is frequently altered in GBM and its deletion or mutation in GBM has been associated with MES transition (9). How PTEN functions upstream of NF-κB pathway to promote MES transition is not well understood in GBM.

The Bromodomain and Extra Terminal (BET) (including BRD2, BRD3, BRD4, and BRDT) family of proteins function as readers for histone lysine acetylation and play a critical role as coactivators in oncogenic transcriptional programs in cancer (10). BRD4 has been shown to bind to acetylated RelA on lysine310 to promote transcription of inflammatory genes in many diseases (11,12). Although BRD2 and BRD4 are overexpressed in GBM (13), their specific role in promoting NF-κB mediated gene expression remains unclear. Due to great interest in the development of BET protein bromodomain specific inhibitors that cross the blood-brain barrier (BBB) (14,15), an improved understanding of the RelA/BET complex and its function in GBM is important to design effective therapeutics.

The NF-κB pathway is under the control of two proteins with opposing functions: PTEN and BET proteins. This interplay implies a potential direct influence of PTEN on the functional modulation of BET proteins within the context of GBM. Here, using patient-derived glioblastoma stem cells (GSCs) and CRISPR gene editing of the RELA gene, we show that RelA is crucial for ECM gene expression and GBM cell invasion associated with MES transition in the context of PTEN loss. Of the BET family proteins, we discovered that BRD2 is dependent on RelA acetylation to be recruited to the promoters of ECM genes and its loss is sufficient to abrogate GBM invasion. Further, we translated our study therapeutically *in vivo* by using a brain penetrant BET-BD2 inhibitor to block mesenchymal gene expression and invasion in mice engrafted with GBM models. These findings reveal a novel mechanism by which loss of PTEN through BRD2 modulates the TME to promote MES transition. Our findings may have important implications for using BD2 specific inhibitors as potent drugs for targeting the MES phenotype in GBM.

## Results

### PTEN negatively regulates chromatin deposition of BRD2 and BRD4 via its phosphatase activity

We previously showed that NF-κB/BRD4-mediated transcription of IL6 and BIRC5 presents a pathway of resistance to tyrosine kinase inhibitors (TKIs) in heterogenous GBM models (16). Here, we investigated the regulation of BRD2 and BRD4 by PTEN as both have been implicated in NF-κB transcription (12). We overexpressed PTEN in patient derived GSCs lacking PTEN expression and assessed chromatin binding of BRD2 and BRD4, which is crucial for their function, by fractionating the nuclear lysate into soluble nuclear extract (SNE) and chromatin bound (CB) fractions. In all GBM cell lines, overexpression of PTEN resulted in the depletion of BRD2 and BRD4 from chromatin, while there was minimal or no change in protein levels observed in SNE fractions (**Figure 1A**). Conversely, knockout of PTEN in a PTEN expressing GBM line (TS576) resulted in an increase in BRD2 and BRD4 on chromatin (**Figure 1B**), suggesting that PTEN regulates the chromatin deposition of these bromodomain proteins in GBM. Phosphatase activity of PTEN is critical for its tumor-suppressive function in multiple cancers (17), including GBM (18,19). To investigate whether the phosphatase activity of PTEN is crucial for its ability to downregulate BRD2 and BRD4 chromatin deposition, we introduced wildtype or a phosphatase dead mutant PTEN (G129R) into U87 cells. Immunoblot analysis revealed an increase in BRD2 and BRD4 levels on chromatin in cells expressing PTEN-G129R compared to those expressing wildtype PTEN (**Figure 1C**), indicating that regulation of BRD2 and BRD4 by PTEN is dependent on its phosphatase activity. PTEN mutations or deletions in GBM and other cancer types lead to the activation of the oncogenic PI3K-AKT pathway (18). To investigate whether the activation of AKT downstream of PTEN loss influences the chromatin binding of BET proteins, we treated U87 cells with an AKT inhibitor and observed decreased chromatin binding of BRD2 and BRD4. These results suggest that AKT activation downstream of PTEN loss regulates the chromatin deposition of BRD2 and BRD4 in GBM (**Figure 1D**).

**Figure 1:**
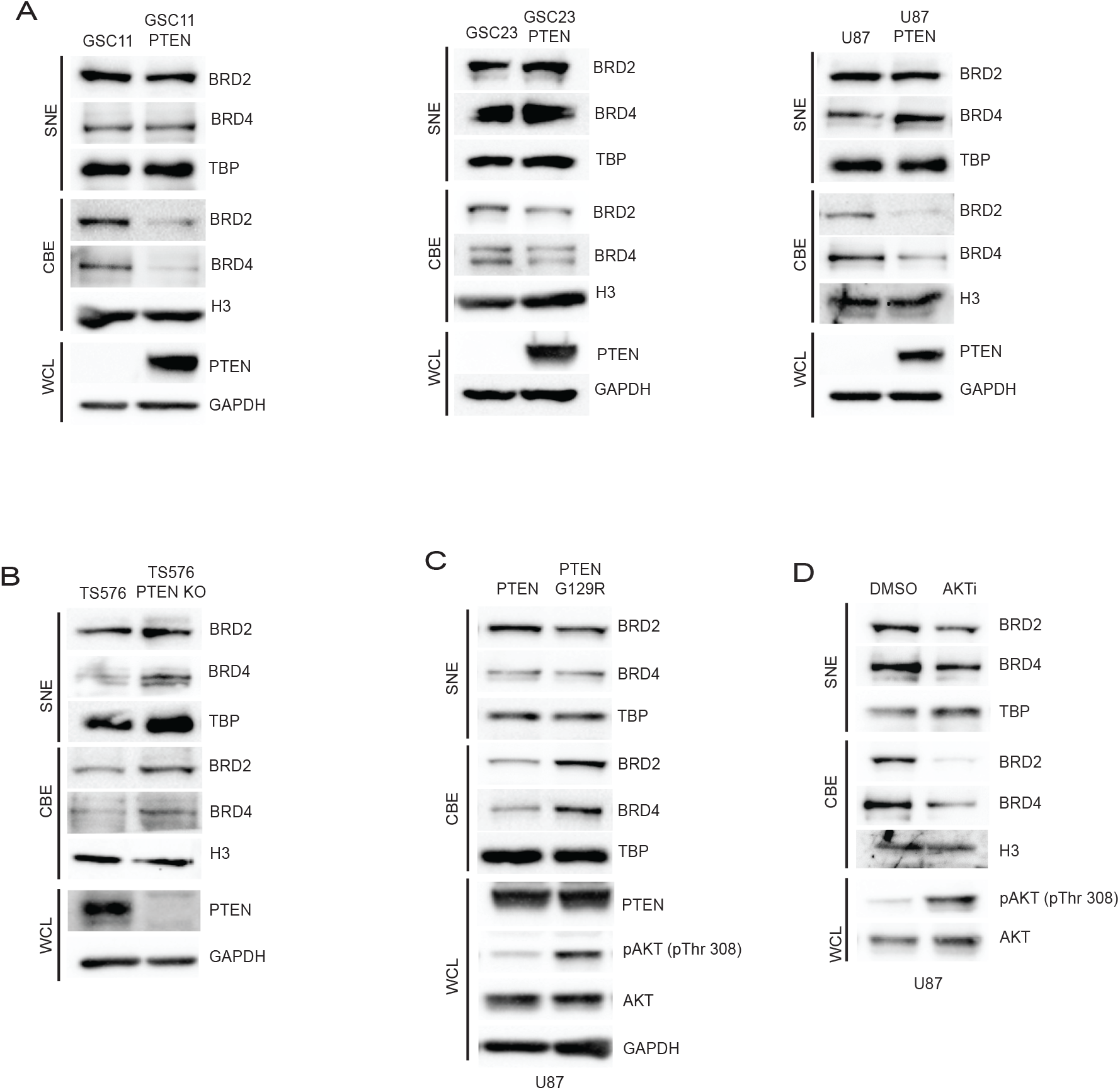
PTEN negatively regulates chromatin deposition of BRD2 and BRD4. Immunoblot analysis of indicated proteins in the soluble nuclear extract (SNE), chromatin bound extract (CBE) and whole cell lysates (WCL) of **A)** GSC11, GSC23 and U87 cells with stably transfected empty vector or PTEN-WT. **B)** TS576 and TS576 PTEN knock out cells. **C)** U87 cells stably expressing wildtype PTEN or phosphatase dead (G129R) PTEN. **D)** GSC11 cells treated with DMSO or Ipatasertib (AKTi, 1uM for 24h). TBP, H3 and GAPDH were used as loading controls for SNE, CBE and WCL extracts respectively.

### Lysine 310 acetylated RelA is crucial for BRD2 and BRD4 chromatin deposition in the absence of PTEN

Our previous data showed the NF-κB complex protein, RelA/p65, recruits BRD4 to promoters of inflammatory genes to facilitate their transcription (16). In GBM, PTEN can regulate the nuclear translocation and transcription of NF-κB (20). Hence, we hypothesized that RelA deposition on chromatin upon PTEN loss may potentially recruit BRD2 and BRD4 to chromatin. To investigate this, we initially evaluated the levels of RelA on chromatin in cells with PTEN deletion and upon PTEN overexpression. Here, we observed increased chromatin associated RelA in PTEN deficient cells, whereas PTEN expression led to downregulation of RelA on chromatin (**Figure 2A**). We also observed increased RelA chromatin binding in cells expressing PTEN-G129R compared to WT-PTEN (**Figure S1A**). Further AKT inhibitor treatment downregulated RelA chromatin levels similar to that observed for BET proteins in Fig.1D (**Figure S1B**). RelA undergoes various post-translational modifications that can impact its stability, subcellular localization, and interactions with other proteins (21). Acetylation of lysine 310 residue on RelA is associated with recruitment of BET proteins to the super enhancers and promoter regions of NF-κB target genes to promote transcription (22). Therefore, we hypothesized that this RelA post-translational modification would facilitate the recruitment of BRD2 and BRD4 proteins to chromatin through their acetylated lysine-binding bromodomains (BD1 and BD2). In support of our hypothesis, we observed an increase in RelA lysine 310 acetylation levels in PTEN deleted cells (**Figure 2B**). To further understand the significance of RelA K310 acetylation in GBM in the context of PTEN loss, we generated an endogenous knock-in at the *RELA* gene by mutating lysine 310 to non-acetylatable arginine (K310R) (**Figure 2C**), here on referred to as RelA-MUT. This mutation caused a modest decrease in RelA protein levels in mutant cells compared to wildtype cells (**Figure 2D**), which might be due to the reported role of K310 acetylation in promoting RelA stability (23). To confirm that K310R mutation does not alter NF-κB function, we assessed RelA-MUT nuclear translocation, activation of a NF-κB reporter, and inhibition by IκBα in response to TNF-α stimulation, which were all confirmed to be equivalent to RelA-WT cells (**Figure 2E** and **S1C**). However, chromatin deposition of BRD2 and BRD4 in RelA-MUT cells was attenuated when compared to RelA-WT cells (**Figure 2F**). In contrast, we did not observe any significant change in the levels of RelA chromatin deposition in mutant cells, suggesting that chromatin bound K310 acetylated protein recruits BRD2 and BRD4 to chromatin in the context of PTEN loss. To identify specific pathway genes that are affected by the disruption of RelA K310 acetylation, we conducted RNA-seq analysis. Surprisingly, the single amino acid change resulted in 8,263 differentially expressed genes in mutant cells when compared to wildtype cells (p_adj_ < 0.05, Benjamini-Hochberg (BH) correction). There were more downregulated (1300, 2-fold decrease, p_adj_ <0.05) than upregulated (668, 2-fold increase, p_adj_ <0.05) genes in RelA MUT cells (**Figure 2G**). Importantly, RelA binding motifs were enriched on the promoters of downregulated genes, suggesting that these genes were under the regulatory control of NF-κB in in complex with BRD2 and/or BRD4 (**Figure S1D**). Gene ontology (GO) analysis performed on downregulated genes in RelA-MUT cells identified biological processes related to extracellular matrix organization (ECM) as genes associated with the top pathway (**Figure 2H**), which overlaps with and is functionally related to the MES gene signature in GBM(24). To confirm that K310 acetylation of RelA is linked to MES gene signature, we assessed CD44 (MES marker) (7) expression and found it downregulated in RelA-MUT cells compared to RelA-WT cells (**Figure S1E**). Further, GSEA enrichment analysis revealed RelA-WT cells were enriched in MES signature whereas RelA-MUT cells were enriched in a PN signature (**Figure 2I**), suggesting that the inhibition of RelA K310 acetylation in the context of PTEN loss could potentially reverse PN to MES transition in GBM. We also assessed the downregulation of MES associated ECM genes (25) by real-time qPCR and observed decreased expression in RelA-MUT cells (**Figure S1F**). Endogenous RelA K310R knock-in mutation in another GSC line (TS576 PTEN KO) displayed similar downregulation of ECM genes by GO analysis (**Figure S1G**). To further support the conclusion that K310 acetylation on RelA is critical for ECM associated gene expression in GBM, we interrogated CBP/P300, known to mediate RelA K310 acetylation (26). Treatment with the CBP/P300 inhibitor, A-485, resulted in a similar enrichment of downregulated ECM genes as observed with RelA-MUT cells (**Figure S1H**), further supporting that acetylation of RelA is a critical regulatory posttranslational modification required for MES gene expression in GBM. This finding is of particular interest since numerous ECM genes have been found to be upregulated in GBM and are associated with a poor prognosis (27).

**Figure 2:**
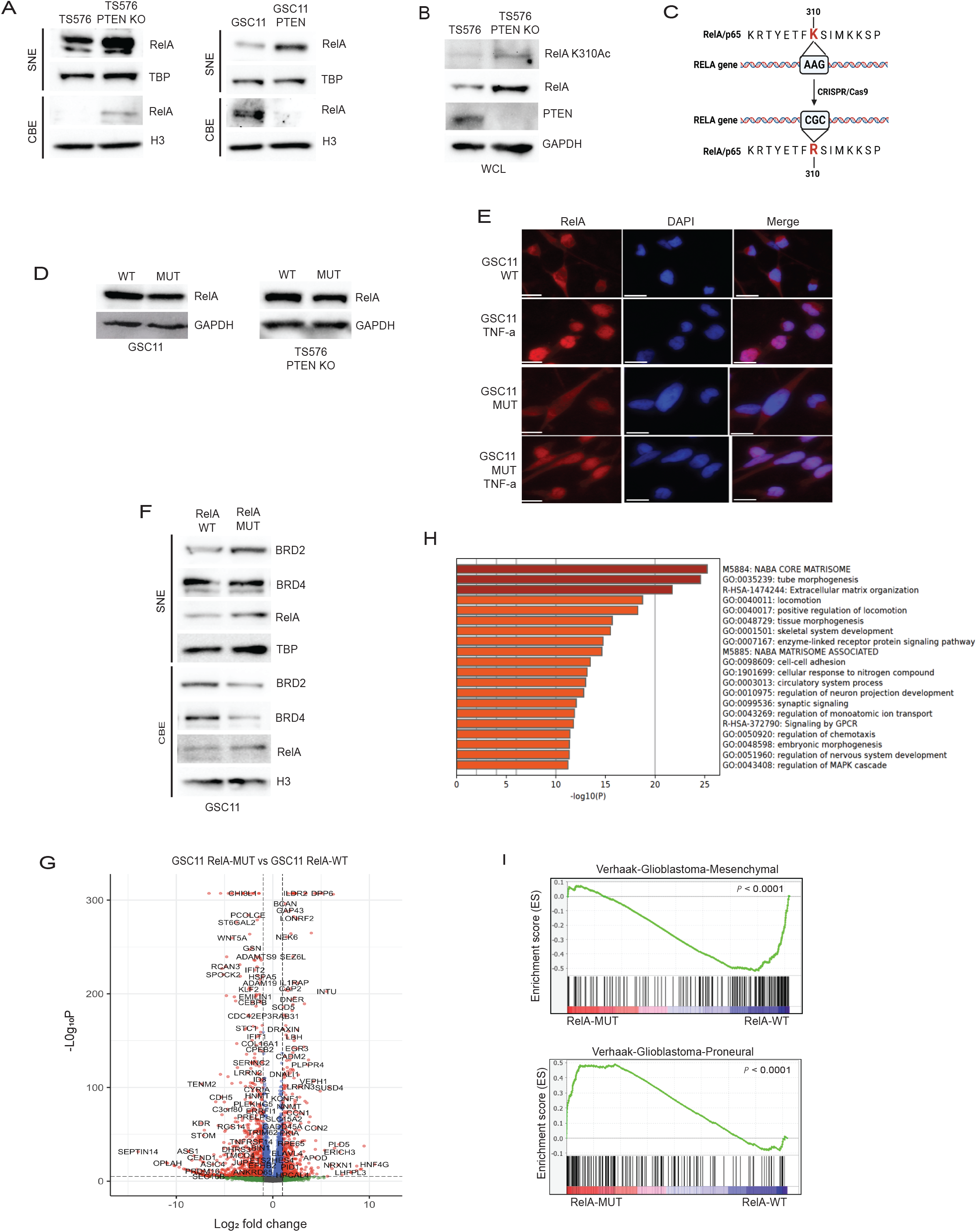
RelA Lysine 310 acetylation recruits BRD2 to BRD4 to chromatin in the absence of PTEN. Immunoblot analysis of **A)** RelA in SNE and CBE fractions of TS576 and GSC11 cells with PTEN deletion or overexpression, respectively; **B)** RelA lysine 310 acetylation in TS576 and TS576 PTEN KO cells. **C)** Schematic for generation of CRISPR knock-in mutation of RELA gene in GBM cell lines. **D)** Immunoblot analysis of RelA in wildtype and mutant GSCs. **E)** Confocal Immunofluorescence images of endogenous RelA nuclear translocation in response to DMSO or TNF-α in GSC11 and GSC11 Rel-MUT cells. Scale bar indicates 20uM. **F)** Immunoblot analysis of indicated proteins in the SNE and CBE fractions of GSC11 RelA-WT and GSC11 RelA-MUT cells. **G)** Volcano plot showing the differential gene expression profiles of GSC11 RelA-WT vs GSC11 RelA-MUT cells. Fold change was plotted as log2(fold change) for each gene relative to its false discovery rate (−log2[FDR]). **H)** GO analysis of genes that are downregulated in RelA mutant cells. **I)** GSEA enrichment plots of GSC11 RelA-WT and GSC11 RelA-MUT gene lists versus queried gene lists are shown. TBP and H3 were used as loading controls for SNE and CBE extracts respectively.

### RelA lysine 310 acetylation promotes BRD2 binding to promoters of ECM genes

To investigate if downregulation of ECM genes in RelA-MUT cells is a result of loss of BET protein recruitment to their promoter regions, we performed ChIP-qPCR assays for RelA, BRD2 and BRD4 co-localization at a series of ECM gene promoters. We observed BRD2 and BRD4 deposition at these promoters (**Figure 3A**) in RelA-WT cells while RelA-MUT ablated BRD2 but not BRD4 deposition, suggesting that BRD2 is dependent on RelA K310 acetylation for recruitment to promoters of ECM genes. Interestingly, we noticed increased RelA-MUT binding at these promoters, which might be due to inefficient removal of RelA from these promoters by IKK due to loss of K310 acetylation (28). However, despite the elevated presence of RelA, we observed a decrease in the RNA expression of the corresponding genes (**Figure S1F**), suggesting BRD2 recruitment to these promoters is crucial for their expression. This dependency on BRD2 deposition to gene regulatory elements occupied by RelA might be important for enhanced transcription of genes where BRD4 is already localized for steady state expression (29). To confirm the role of BRD2 for ECM gene expression, we depleted BRD2 by shRNA knockdown and assessed gene expression by RT-qPCR (**Figure 3B**), which further illustrated a BRD2 dependent ECM gene signature. Furthermore, RNA-seq analysis indicated a greater number of genes were downregulated in cells with BRD2 knockdown as compared to control cells (**Figure S2A**), with GO analysis revealing genes related to MES phenotype, such as ECM, migration and wound healing, being predominantly affected (**Figure 3C**). We also determined overlapping genes affected by RelA-MUT and BRD2 knockdown and found 213 genes shared in these datasets (**Figure 3D**). GO analysis of this intersection revealed ECM associated genes were the top downregulated genes (**Figure 3E**). This finding provides additional evidence supporting the regulatory role of RelA/BRD2 in controlling the expression of genes associated with MES phenotype. Further, we performed GSEA and observed an enrichment of the PN signature in BRD2 knockdown cells, whereas control shRNA cells displayed an enrichment of a MES signature (**Figure 3F**). Taken together, these findings suggest that BRD2 is a crucial chromatin regulator for MES gene expression in GBM. Consequently, its loss leads to an enrichment of the PN phenotype, underscoring the dynamic nature of cellular states in GBM and the pivotal role of RelA/BRD2 complex in these transitions.

**Figure 3:**
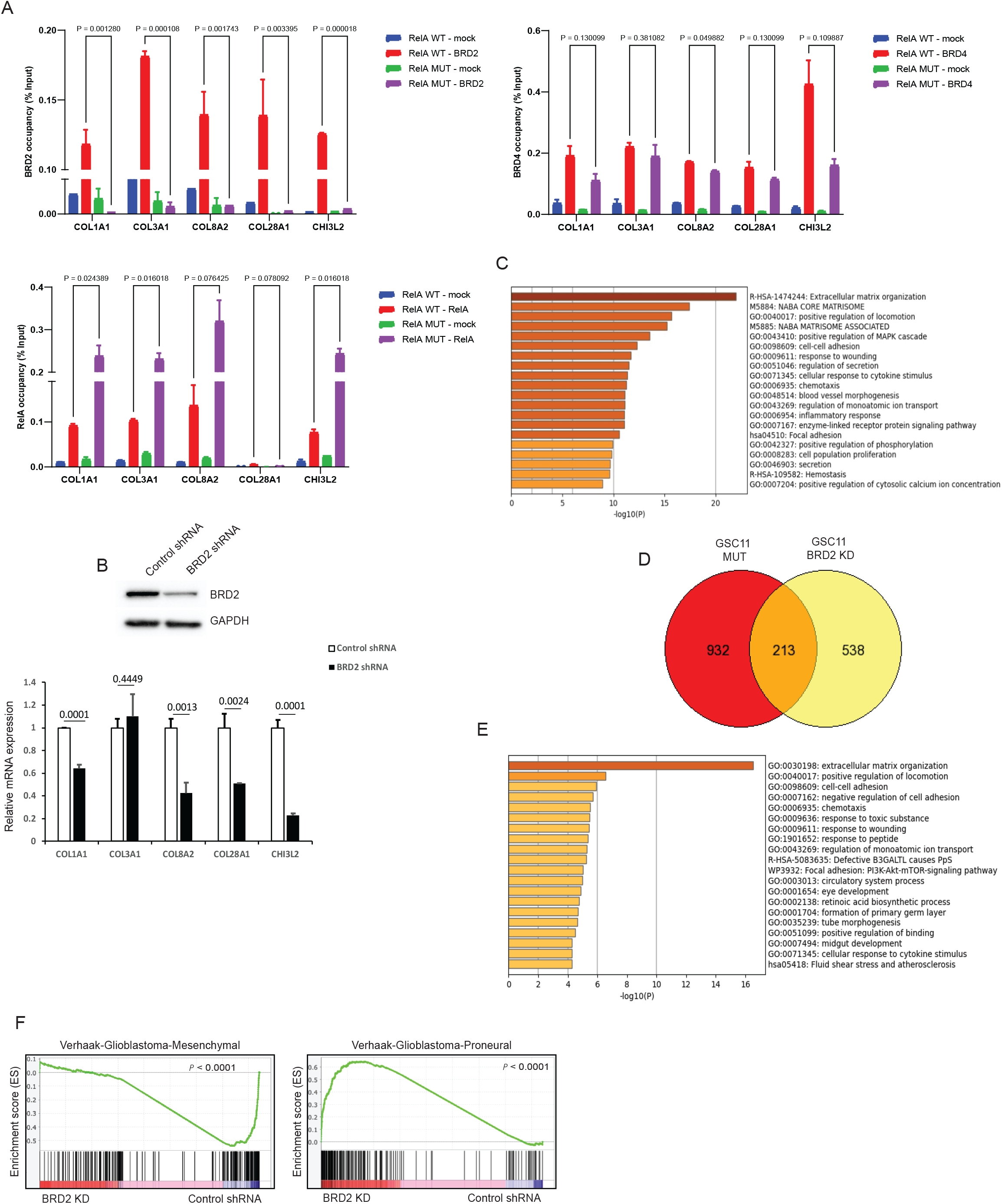
Expression of ECM genes is dependent on BRD2 chromatin binding but not BRD4. **A)** ChIP-qPCR with anti-BRD2, BRD4 and RelA on the promoters of ECM genes in GSC11 RelA-WT and GSC11 RelA-MUT cells. For ChIP assays, bar graphs indicate fold enrichment of BRD2, BRD4 and RelA over the input (n=3 biological samples with three replicates each). Data represents mean ± SD. **B**) Immunoblot analysis of BRD2 expression in control shRNA or BRD2 shRNA expressing GSC11 cells (top panel). RT-qPCR analysis of ECM genes in control shRNA or BRD2 shRNA expressing GSC11 cells. Data represents mean ± SD. **C)** GO analysis of genes that are downregulated in BRD2 shRNA expressing cells. **D)** Venn diagram showing shared genes that are downregulated in GSC11 RelA-MUT cells and GSC11 BRD2 knockdown cells. **E)** GO analysis of shared downregulated genes in Fig.3D**. F)** GSEA enrichment plots of GSC11 BRD2 shRNA and GSC11 control shRNA gene lists versus queried gene lists are shown.

### Decreased invasion of RelA mutant cells is phenocopied by BRD2 downregulation

ECM plays an important role in promoting the migration of glioma cells (29). Invasive GSCs associated with MES subtype express high ECM gene expression and have been linked to poor GBM patient survival (24). We sought to evaluate the invasive properties of the RelA mutant GSCs. Accordingly, we performed a matrigel invasion assay to compare the migration of RelA-MUT and RelA-WT GSCs. RelA-MUT GSCs exhibited significantly reduced invasion compared to parental cells in response to growth factors (**Figure 4A**). *In vitro* growth assays revealed GSC11 RelA-MUT cells displayed slower growth compared to wildtype cells (**Figure S3A**). To evaluate the tumor forming capacity of these mutant GSCs in vivo, we orthotopically engrafted RelA-WT or RelA-MUT cell lines into mice. Surprisingly, RelA mutant GSC11 failed to initiate tumor growth in vivo (data not shown). In contrast, GSC TS576 with *PTEN* KO and RelA K310R mutation exhibited comparable *in vitro* growth rates when compared to RelA wildtype cells (**Figure S3B**). The variation in cell growth properties observed in RelA-MUT cells could potentially stem from differences in their genomic alterations. Although the isogenic TS576 lines had comparable *in vitro* growth rates, RelA-MUT cells had notably smaller tumors than RelA-WT cells upon orthotopic engraftment (**Figure 4B**). These differences in proliferation within an *in vivo* environment may be due to an interaction between the RelA-MUT cancer cells and the microenvironment. Tumor associated macrophages/microglia (TAMs) have been associated with promoting tumor invasion in GBM (30). To determine if reduced invasion and growth of RelA-MUT cells resulted from a decrease in TAM infiltration, we conducted immunohistochemistry (IHC) analysis on tumor sections. IHC revealed a higher number of IBA1^+^ invasive GBM cells at the tumor periphery, accompanied by elevated levels of TAMs in RelA-WT cells compared to RelA-MUT cells (**Figure 4C**). Subsequently, we evaluated if the depletion of BRD2 phenocopies RelA K310R mutation and observed reduced invasion of BRD2 knockdown cells compared to control shRNA cells (**Figure 4D and S3C),** with a modest effect on the proliferation of TS576 *PTEN* KO cells (**Figure S3D-E**). Since the MES gene signature is associated with therapeutic resistance in GBM (31), we assessed whether depletion of BRD2 sensitizes GSCs to temozolomide (TMZ) and found that BRD2 knockdown cells were more sensitive to TMZ compared to control GSCs (**Figure 4E and S3F**). Overall, these data suggests that RelA/BRD2 promotes a MES phenotype in GBM.

**Figure 4:**
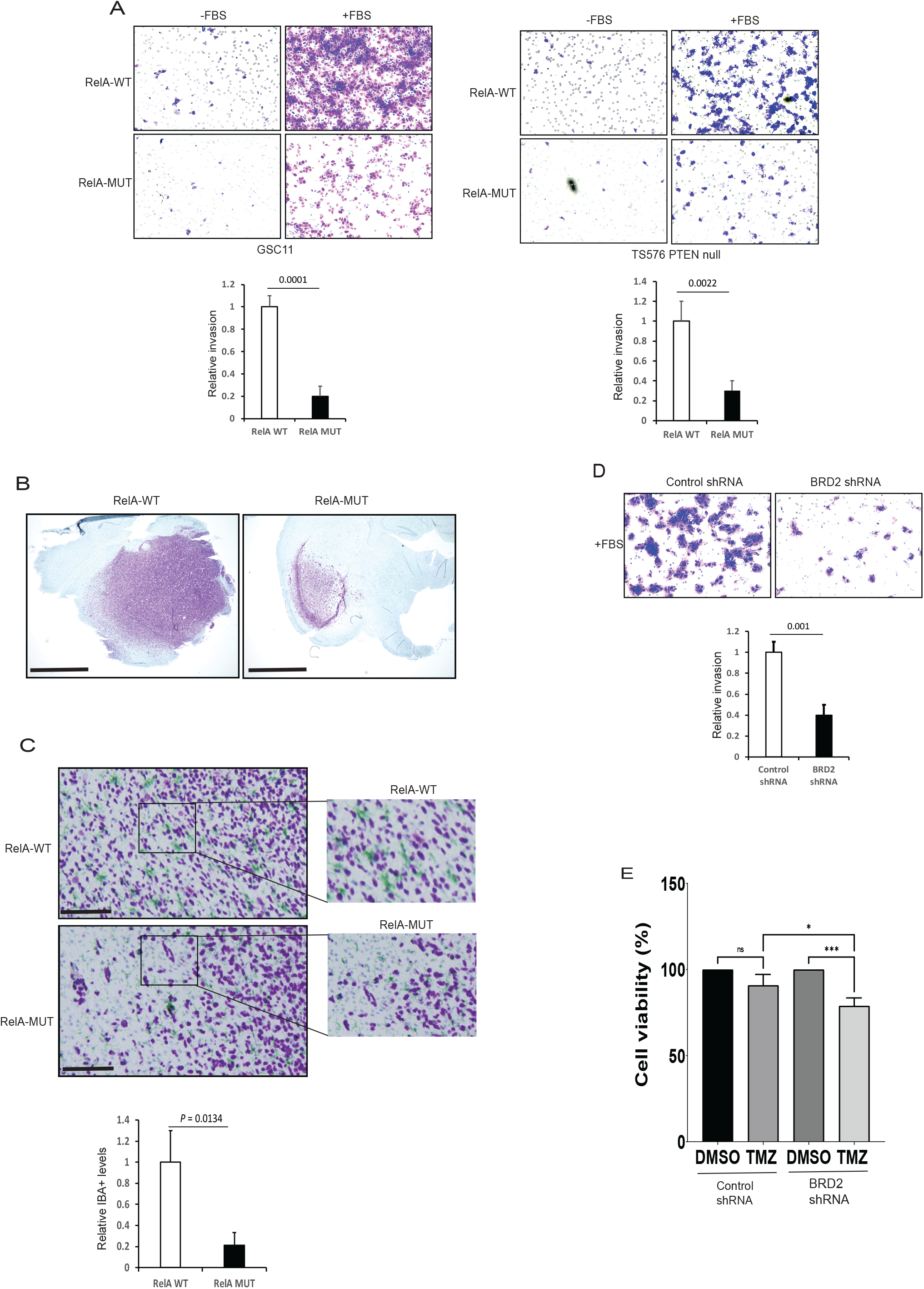
MES phenotype is dependent on K310 acetylated RelA recruitment of BRD2 to chromatin. **A)** Matrigel cell invasion of RelA-WT and RelA-MUT cells were examined by trans well-chamber assays. The number of invading tumor cells that penetrated through the Matrigel was counted using 5 randomly selected fields and expressed as relative percentage. Data represents mean ± SD. **B)** Immunostaining showing *in vivo* growth of RelA-WT and RelA-MUT GSCs. Glioma cells (NM95; purple) **C)** Immunostaining showing *in vivo* invasion of RelA-WT and RelA-MUT GSCs. Glioma cells (NM95; purple) and macrophages/microglia (IBA1; green). Enlarged images of Figure C with quantification of IBA+ cells surrounding GBM cells. **D)** Matrigel invasion assay of GSC11 control shRNA or BRD2 shRNA expressing cells. Quantification was performed as in A. **E)** Control or BRD2 expressing GSC11 cells were treated with 100uM TMZ for 24 h and cell viability was assessed by ATPlite assay. Values represent mean of 3 experiments ± SD. All Data represents mean ± SD.

### BET-BD2 specific inhibition downregulates ECM gene expression and attenuates invasion of GSCs

Pan-BET family inhibitors have been shown to have significant toxicity in clinical studies (32). In contrast, BD2-specific inhibitors, selectively targeting the second bromodomain of BET proteins, have been found to be less toxic in preclinical studies (29,33). BRD4 has been associated with ECM expression in diverse fibrotic diseases (34). Based on this finding, a recently developed BET-BD2 inhibitor has advanced to clinical trials for the treatment of myelofibrosis (NCT04454658). Therefore, we hypothesized that BD2 specific inhibition might be effective in downregulating ECM expression and MES signature in GBM with less toxicity. To initially assess whether a BD2 inhibitor (BD2i) could displace BET family proteins from chromatin, we treated cells with GSK620 (29) and analyzed chromatin bound fractions for BET family proteins and observed reduced chromatin deposition of BRD2, but not BRD3 or BRD4, when compared to vehicle treatment (**Figure 5A**). Chromatin bound RelA levels remained unchanged with BD2i treatment indicating drug specificity (**Figure 5A**). Next, to explore genes that were differentially regulated in BD2i treated cells, we conducted RNA-seq analysis and found a total of 3,048 significantly differentially expressed genes (p_ad_j < 0.05, **Figure 5B**). Furthermore, GO analysis of downregulated genes revealed ECM pathway genes were the top differentially expressed genes (**Figure 5C**). Importantly, RelA regulatory elements were enriched on the promoters of downregulated genes, suggesting that these genes were under the regulatory control of NF-κB (**Figure S4A**). A second GSC line displayed similar downregulation of ECM genes by GO analysis (**Figure S4B**). To determine genes similarly affected by RelA-MUT and by BD2i treatment, we analyzed the intersection of both conditions and identified approximately 50% of the genes downregulated in BD2i treatment were shared with RelA-MUT cells (127 genes total) (**Figure 5D**). GO analysis of shared genes revealed ECM and multiple oncogenic pathways associated with GBM, such as angiogenesis and PI3K signaling, were also enriched across shared significantly decreased genes (**Figure 5E**). Furthermore, RT-qPCR analysis confirmed downregulation of many MES genes with BD2i treatment (**Figure 5F**). A second BD2i, ABBV-774(33) similarly downregulated MES gene expression (**Figure S4C**). GSEA enrichment analysis revealed control cells were enriched in MES signature compared to BD2i treated cells (**Figure 5G**), suggesting that ECM gene downregulation led to loss of MES transition. Interestingly, unlike other perturbation conditions, BD2i treated cells are not enriched in the proneural transcriptional subtype despite the loss of mesenchymal gene expression (**Figure S4D**). To confirm whether BRD2 levels on the promoters of ECM genes were downregulated with BD2i treatment, we performed ChIP-qPCR and observed significant BRD2 depletion from chromatin (**Figure 5H**). In contrast, we did not observe any changes in BRD4 levels on these promoters, suggesting that GSK620 specifically displaces BRD2 from these promoters (**Figure S4E**). Furthermore, by matrigel invasion assay we detected attenuated invasion for BD2i treated cells when compared to DMSO (**Figure 5I and S4F**).

**Figure 5.**
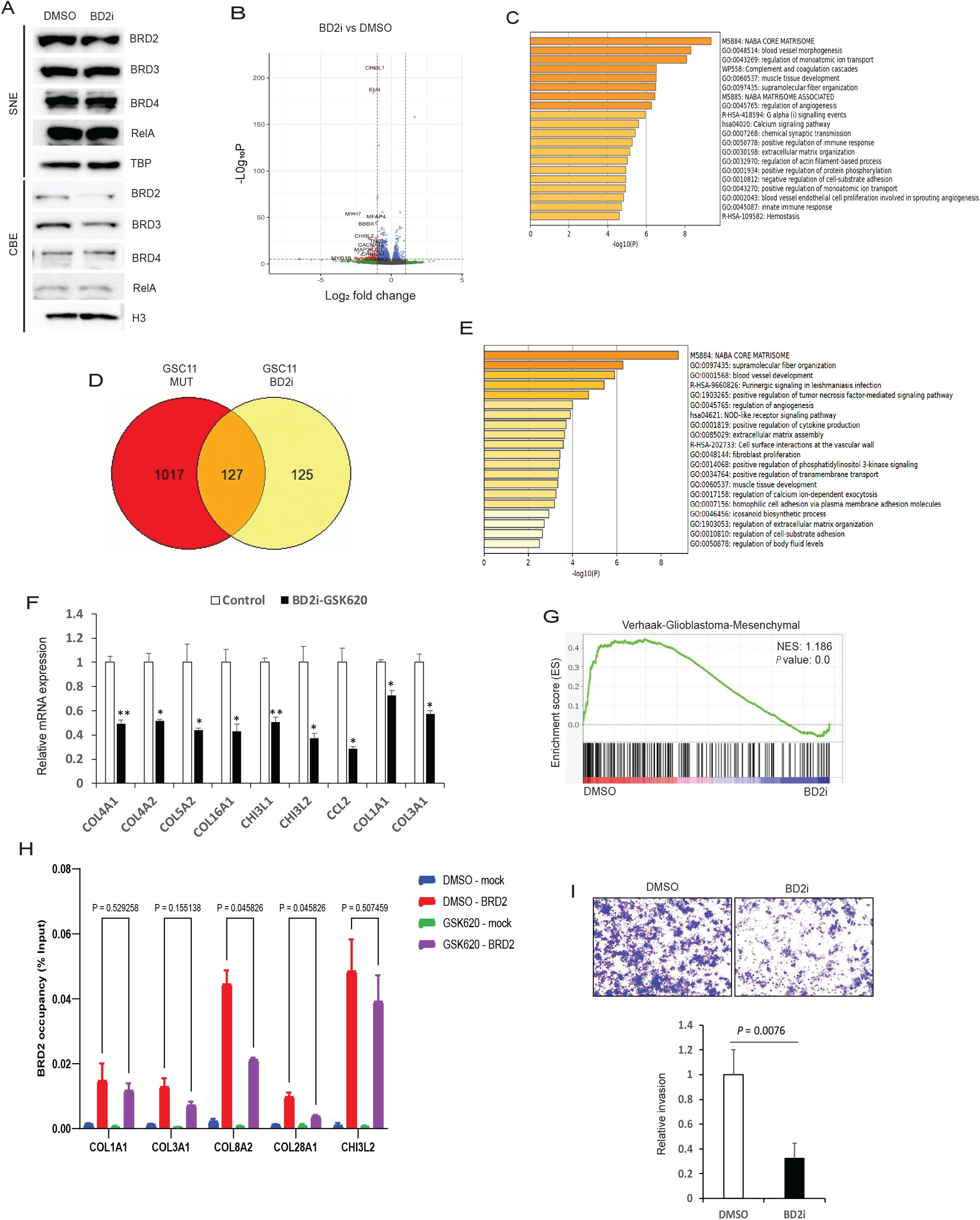
BET-BD2 inhibitors mimic RelA K310R mutant in regulating ECM genes expression and invasion. **A)** Immunoblot analysis of indicated proteins in SNE and CBE fractions of GSC11 cells treated with DMSO or 0.5uM of GSK620 (BD2i) for 24h. **B)** Volcano plot showing the differential gene expression profiles of DMSO or BD2i (0.5 uM/24h) treated GSC11 cells. Fold change was plotted as log2(fold change) for each gene relative to its false discovery rate (−log2[FDR]). **C)** GO analysis of genes that are downregulated in BD2i treated cells compared to DMSO treated GSC11 cells. **D)** Venn diagram showing shared genes downregulated in GSC11 RelA-MUT cells and BD2i treated GSC11 cells. **E)** GO analysis of shared downregulated genes in Fig.5D. **F)** RT-qPCR analysis of mesenchymal genes in BD2i treated cells. GSEA enrichment plots of DMSO or BD2i gene lists versus MES gene expression is shown. **G)** GSEA enrichment enrichment plots of DMSO or BD2i gene lists versus MES gene expression is shown. **H)** ChIP-qPCR with anti-BRD2 on the promoters of ECM genes in DMSO and BD2i treated GSC11 cells. For ChIP assays, bar graphs indicate fold enrichment of BRD2 over the input (n=3 biological samples with three replicates each). **I)** Matrigel cell invasion of GSC11 cells treated with DMSO or BD2i for 24h were examined by transwell-chamber assays. The number of invading tumor cells that penetrated through the Matrigel was counted using 5 randomly selected fields and expressed as relative percentage. Values represent the means of 3 experiments ± SD.

### Treatment with BET-BD2 inhibitor reduces MES phenotype *in vivo*

We evaluated the effect of GSK620 on cell proliferation by treating GSCs with increasing doses of GSK620. We observed GSC11 cells displayed decreased cell proliferation at 1 μM dose (**Figure 6A**). However, another cell line GSC23, did not exhibit any changes in cell proliferation even at higher doses of GSK620 (**Figure S5A**), suggesting the differential sensitivities of GSCs to BD2i *in vitro*. Subsequently, we explored whether GSK620 could enhance the sensitivity of the resistant GSC23 cells to ionizing radiation (IR). We pre-treated GSC23 cells for 24h with vehicle or GSK620 and exposed to one dose of IR (4Gy) and assessed the growth of these cells for a period of 5 days. We observed a significant reduction in the proliferation of GSK620-treated GSC23 cells subjected to IR compared to the control group that received only IR treatment (**Figure 6B**). This suggests that the inhibition of MES gene expression in these cells with BD2i could have increased their sensitivity to IR. Next, to determine the ability of GSK620 to traverse the BBB and hence its candidacy for *in vivo* use, pharmacokinetic (PK) analysis was performed on mice administered GSK620 by oral gavage. Analysis of the brain/plasma ratio revealed an AUC_0-24hr_ of 27.2% and plasma t1/2 of 1.05 hrs (**Figure S5B**) which is comparable to TMZ kinetics (35), suggesting that GSK620 has adequate CNS penetration. Next, to assess whether targeting the ECM gene expression in GBM models *in vivo* leads to loss of the MES phenotype, we orthotopically engrafted tumor cells into mice. Following tumor establishment, as assessed by near infrared imaging (36), mice were treated with vehicle or GSK620 for alternative days and monitored for tumor burden. Tumors from moribund mice were assessed for invasion and TAMs, both of which are associated with the MES phenotype (37) (**Figure 6C**). We did not detect any differences in tumor volume between vehicle and BD2i treated mice (**Figure 6D**). IHC analysis of sectioned brains showed tumors with diffuse infiltration into the surrounding brain parenchyma for vehicle treated animals (**Figure 6E**), while BD2i treatment was associated with less diffuse and more localized tumor growth (**Figure 6E**). Gliomas invade most notably along white matter tracts, perivascular spaces, and meninges, which stand out as key clinicopathological features of these aggressive tumors (38). We assessed the corpus callosum region which contained invading cells into the contralateral hemisphere (**Figure 6F**) (38). We also observed that MES-like invasive regions with closely aligned bundled cells with TAMs were significantly reduced in GSK620 treated mice compared to vehicle (**Figure 6F**), suggesting that BD2i inhibited a MES like phenotypic growth pattern with high TAMs *in vivo*. Lastly, to assess whether our findings of PTEN regulation of BET proteins levels *in vitro* are also observed in patient GBM samples, we analyzed recently published GBM patient proteomic data (39). Interestingly, our Pearson correlation analysis displayed a significant inverse correlation between PTEN and BRD2 (P = 0.0023) but not with BRD3 and BRD4 proteins in GBM patients (**Figure S5C**). These results support our conclusion that BRD2 is a key effector BET protein to promote MES transition downstream of PTEN loss in GBM (**Figure 7**).

**Figure 6:**
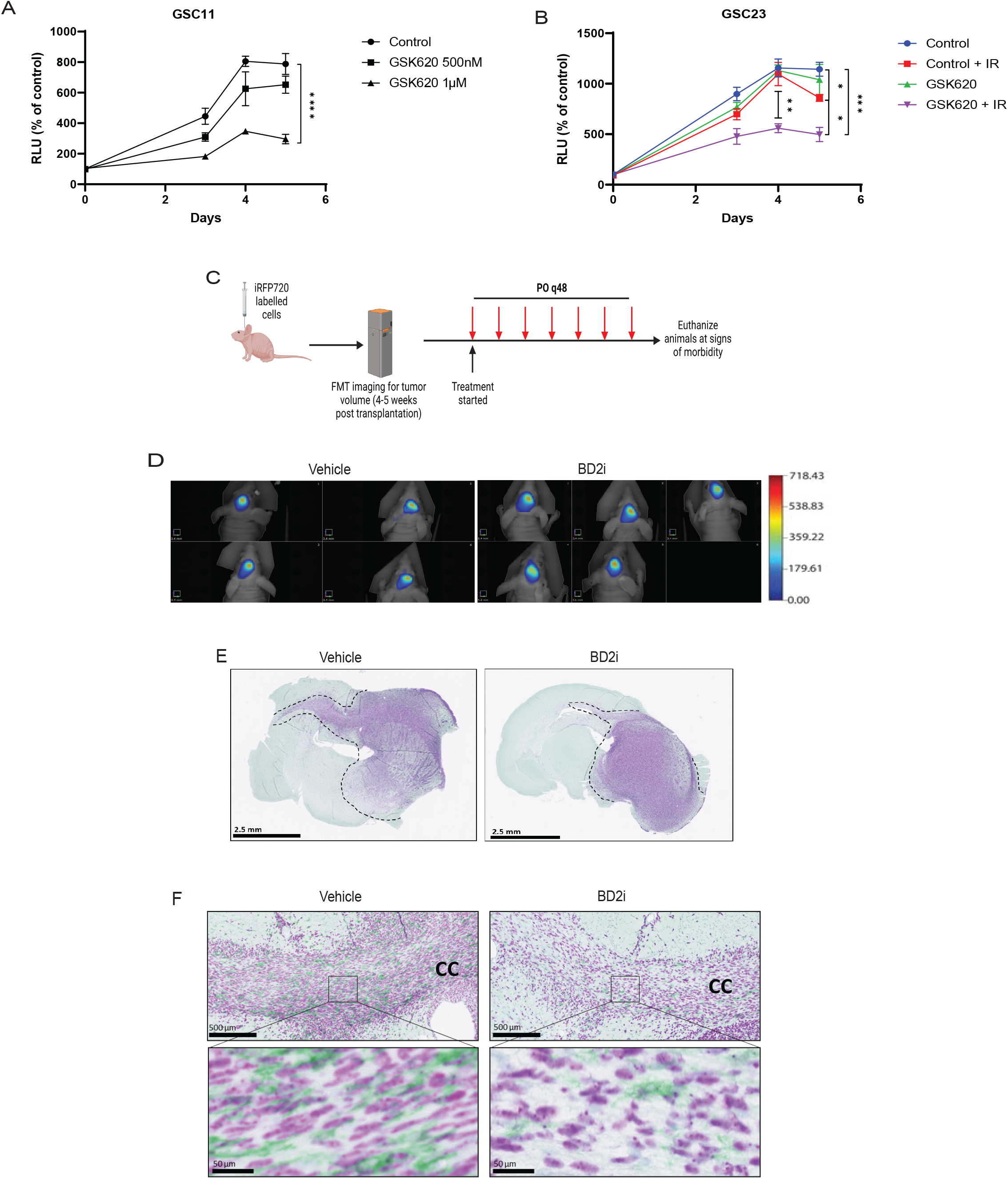
BD2i treatment attenuates glioma invasion *in vivo*. **A)** Effect of GSK620 on GSC11 *in vitro* cell growth by ATPlite assay. **B)** GSC23 cells pre-treated with DMSO or GSK620 (2 μM) for 24h were exposed to one dose of IR (4Gy) and assessed for cell growth for 5 days by ATPlite assay. **C)** Experimental drug trial design: Mice were orthotopically transplanted with iRFP-720-labelled GSC11 cells (1 x 10^5^ cells). Tumor burden was assessed by FMT imaging and mice were randomly assigned to vehicle or GSK620 treatment. **D)** FMT images showing tumor burden in vehicle or GSK620 treated mice. n = 5. **E)** Immunostaining showing invasive growth of GSC11 tumors in mice treated with vehicle or BD2i (NM95; purple for human nuclei). **F)** Immunostaining showing invading cells along the white matter tracts (corpus callosum) with TAMs in vehicle or BD2i treated mice (NM95; IBA1; green for TAMs).

**Figure 7:**
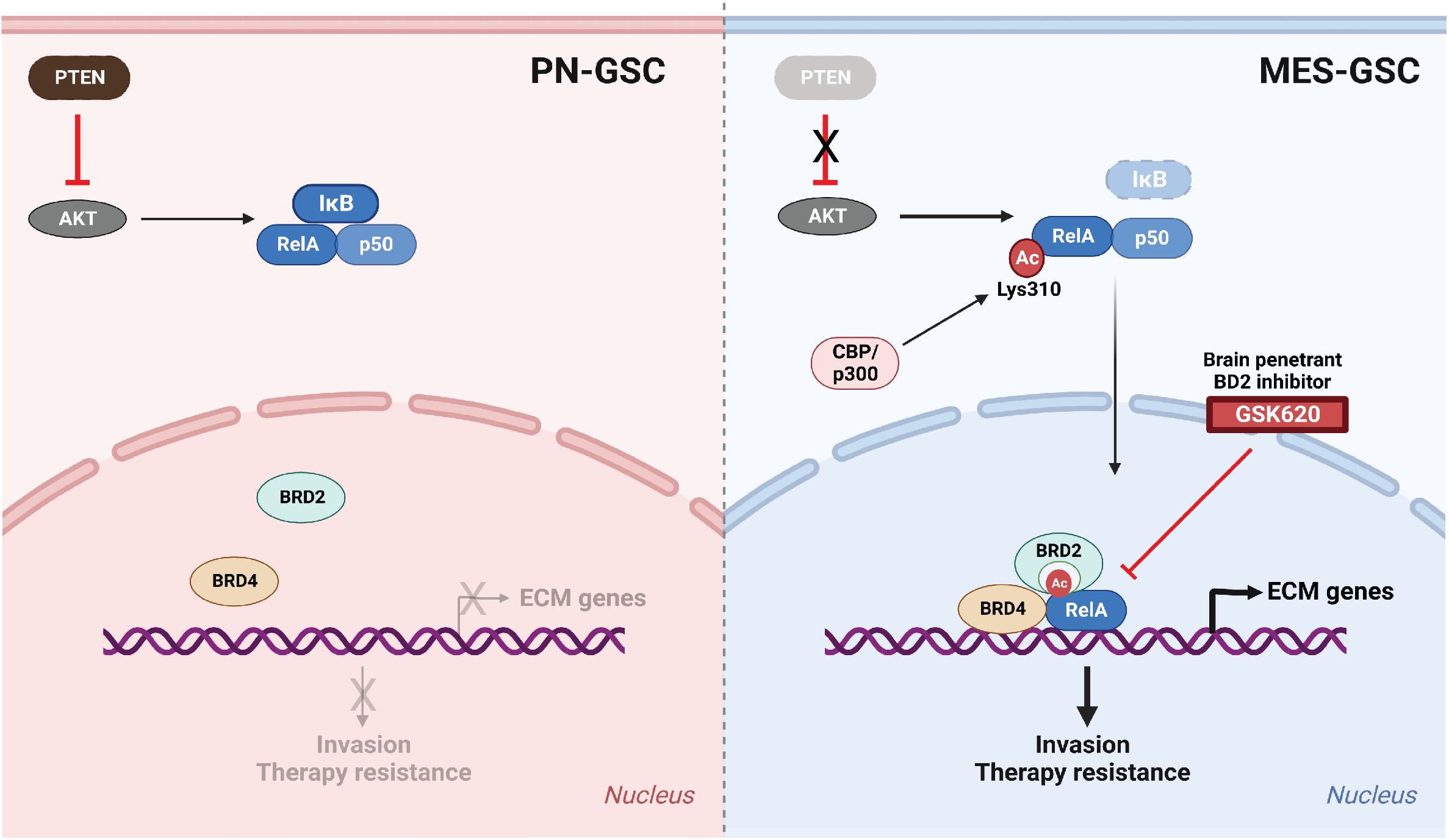
Schematic of PTEN-NFκB-BRD2 axis in driving mesenchymal transition in GBM

## Discussion

GBM is a highly aggressive brain tumor characterized by its infiltrative nature and resistance to conventional therapies. The TME plays a crucial role in GBM pathogenesis, with the interaction between cellular and non-cellular components being a major determinant of treatment outcomes (40). ECM, a non-cellular component constitutes a complex structural network that provides support and facilitates cellular growth, survival, maturation, differentiation, and migration. Here, we report the role of the PTEN-NF-κB-BRD2 axis in regulating ECM gene expression and MES transition leading to invasiveness of GBM cells. Our findings revealed that NF-κB is a key player downstream of PTEN loss in promoting ECM gene expression by regulating the recruitment of BRD2 to the promoters of these genes through its lysine 310 acetylation (**Figure 7**).

Dysregulation of ECM genes in GBM promotes the infiltration of tumor cells into the surrounding brain parenchyma, resulting in nonresectable tumors that subsequently lead to tumor recurrence (1,27). Recent findings on ECM gene regulation in GBM underscore the significance of targeting ECM components as a potential therapeutic approach for treating this malignancy (25,27,41). Previous research has shown that in the context of pulmonary fibrosis, BRD4 regulates the expression of NF-κB dependent EMT genes (42), suggesting that a similar mechanism could be at play in tumors as well. Nonetheless, a recent study utilizing a CRISPR screen found BRD2, rather than BRD3 or BRD4, promotes EMT in lung cancer models (43). Aligning with these findings, our data show that BRD2 and BRD4 are localized at the promoters of NF-κB targeted mesenchymal genes. Remarkably, we found that mere downregulation of BRD2 is sufficient to attenuate expression of ECM genes and invasion in GBM.

Recent studies highlighted the role of targeting ECM in GBM. One study showed that invasive regions associated with MES like cells, oncostreams, are enriched with ECM genes. This study also showed that targeting the COL1A1 protein led to loss of oncostream characteristics and prolonged survival in mouse GBM models (44). Another study reported the association of MES transition with ECM gene expression with a negative prognostic value at recurrence, suggesting a link between ECM expression and invasion with therapy resistance in recurrent GBM (45). We demonstrate that the inhibition of BRD2 deposition on ECM gene promoters by RELA K310R mutation, knockdown or using BET-BD2-specific inhibitors led to loss of ECM gene expression, MES transition and invasion of GBM cells, ultimately resulting in decreased macrophage infiltration in *in vivo* GBM models. This is particularly significant given the association between PTEN deletion and increased TAM infiltration, which contributes to the mesenchymal phenotype in GBM (9,46). Our study also highlights the need to define how BRD2 regulate chromatin and transcription in processes such as cell-fate transitions and reprogramming in GBM, since BRD2 has been shown to play a role in chromatin organization (47,48) and its mutations have been associated with other CNS disorders (49).

Although it is widely accepted that NF-κB is aberrantly activated in GBM and drives MES transition, direct targeting of NF-κB has been challenging due to the lack of inhibitors that selectively block its function without affecting other crucial cellular pathways. Our data reveals a previously unknown mechanism in which acetylation of RelA at K310 modulates the MES transition by regulating ECM genes through BET proteins. Intriguingly, we observed that RelA-MUT GSC11, a recurrent PDX model, loses MES expression and gains PN expression, suggesting that the MES transition is linked to epigenetic alterations as seen in other tumor types (50). These RelA-MUT GSC11 lines failed to initiate tumor growth *in vivo,* highlighting its reliance on NF-κB signaling. Another PDX line, TS576 engineered with *PTEN* knockout made significantly smaller tumors *in vivo* in the context of the RelA K310R mutation. These differences between the cell lines could be due to different additional genomic alterations in these PDX lines; however, invasion and ECM gene expression were the core common pathways downregulated between these two lines, suggesting the acetylation of RelA/p65 plays a key regulatory switch in these GBM processes.

We did not find any survival benefit using BET-BD2 inhibitor alone in PDX models. Since MES tumors are resistant to therapy (31), we anticipate combining BD2i with chemo or radiation therapy may result in a survival benefit. Indeed, our *in vitro* experiments showed that BRD2 knockdown cells become more responsive to TMZ and BD2i sensitized GSCs to IR therapy.

Overall, our study highlights the potential of targeting the PTEN-NF-κB-BRD2 axis using BET-BD2 inhibitors as a novel therapeutic strategy for targeting MES transition in GBM. This approach may be especially effective in tumors with non-functional PTEN, as it might overcome the limitations of immune checkpoint inhibitors and enhance anti-tumor immunity by modulating the immunosuppressive TME (29,33). Further research is warranted to evaluate the safety and efficacy of BET-BD2 inhibitors in GBM patients with inactivated PTEN and to optimize treatment strategies for this devastating disease.

## Materials and methods

### Cell culture

Parental U87MG were obtained and cultured as described previously (51,52). PDX sphere lines were cultured in DMEM/F12 medium supplemented with B27 (GIBCO/Life Technologies) and 20 ng/ mL human recombinant EGF, 20 ng/mL bFGF. GSC11 was provided by Frederick Lang (M.D. Anderson Cancer Center); TS576 was provided by Cameron Brennan (Memorial Sloan Kettering Cancer Center). All cells were incubated at 37°C, 5% CO2, and 100% relative humidity in low-attachment flasks. PDX cell lines were dissociated with Accutase (Stemcell Technologies). GSK620 and A-485 were purchased from MedChem Express. ABBV-744 was provided by Andrew Shiau (UCSD). The near-infrared fluorescent protein iRFP720 cDNA construct was from (53). pLV-IκB-SR vector was a gift from Inder Verma (Salk Institute). The pGL4.32[luc2P/ NF-κB-RE/Hygro] vector for NF-κB luciferase reporter assays was purchased from Promega. shRNA constructs targeting BRD2 were purchased from Sigma (Mission shRNA).

### Protein subcellular fractionation and western blotting

Cultured cells were pelleted, and the supernatant was discarded, leaving the cells as dry as possible. Thermo Scientific Subcellular Protein Fractionation Kit for Cultured Cells was employed to separate different protein cell compartments. Cytoplasmic, nuclear and chromatin-bound proteins were extracted according to manufacturer’s instructions (#78840). For whole cell lysates, cells were lysed in RIPA buffer. Extracts were separated using gel electrophoresis and transferred via wet transfer onto a PVDF membrane. The membrane was blocked with 5% milk in TBST and probed with primary antibodies in 5% BSA at 1:1,000 dilution overnight at 4°C and secondary HRP antibodies in 5% milk at 1:10,000 for 1 hour at RT. Signal was assessed via chemiluminescence with the SuperSignal West Pico PLUS substrate (Thermo Fisher, #34580) and visualized on a ChemiDoc MP system (Bio-Rad). The following antibodies were used, anti-BRD2 (Cell Signaling Technology, #5848), anti-RelA/p65 (Cell Signaling Technology, #8242), anti-BRD4 (Cell Signaling Technology, #13440), anti-GAPDH (Cell Signaling Technology, #2118), anti-TBP (Cell Signaling Technology, #44059), PTEN (Milliopore, #04-035), H3 (Novus Biologicals, #NBP1-61519), AKT (Cell Signaling Technology, #9272), anti-phospho-AKT Thr308 (Cell Signaling Technology, #9275), anti-acetyl-NF-κB p65 (Cell Signaling Technology, #3045), anti-BRD3 (Santa Cruz Biotechnology, #sc-81202).

### Generation of CRISPR engineered cell lines

pSpCas9(BB)-2A-GFP (px458) plasmid was a gift from Feng Zhang (Addgene plasmid #48138). The designated sgRNA sequences for each of the targeted genes were cloned into px458 using combinations of top and bottom oligonucleotides listed below.

PTEN – guide 1-top: 5’ – CACCGGAATTTACGCTATACGGAC – 3’

PTEN – guide 1-bottom: 5’ – AAACGTCCGTATAGCGTAAATTCCC -3’

PTEN – guide 2-top: 5’ – CACCGAACAAGATCTGAAGCTCTAC – 3’

PTEN – guide 2-bottom: 5’ – AAACGTAGAGCTTCAGATCTTGTTC – 3’

RELA-K310R-top: 5’ – CACCGCTTCTTCATGATGCTCTTGA – 3’

RELA-K310R-bottom: 5’ – AAACTCAAGAGCATCATGAAGAAGC – 3’

Each pair of top and bottom oligonucleotides were phosphorylated and annealed by incubating 10 µM of each with 1 × T4 DNA ligase buffer (New England Biolabs), 5U T4 polynucleotide kinase (New England Biolabs) at 37 °C for 30 min, 95 °C for 5 min and by cooling down to 25 °C at 0.1 °C/s using a thermocycler. Annealed oligonucleotides were cloned into px458 by incubating 25 ng px458, 1 μM annealed oligonucleotides, 1× CutSmart buffer (New England Biolabs), 1 mM ATP (New England Biolabs), 10U BBSI-HF (New England Biolabs) and 200U T4 ligase (New England Biolabs) at 37 °C for 5 minutes, 23 °C for 5 min for 30 cycles. Correct cloning of each sgRNA sequence was confirmed by Sanger sequencing using U6 sequencing primer: 5′-GATACAAGGCTGTTAGAGAGATAATT-3′.

A single-stranded oligo DNA nucleotides (ssODNs) listed below was used to introduce the point mutation into the RELA gene.

RELA-K310R-ssODN

5’ - CCTTACTTTCCCAGACGATCGTCACCGGATTGAGGAGAAACGTAAAAGGACATATGAGACATTCCGCAGCATCATGAAGAAGAGTCCTTTCAGCGGTGAGATGGGGACTGGGAAAGCCAGAGAGGAA - 3’

GSCs were dissociated to single cells using Accutase (Innovative Cell Technologies). The dissociated GSCs (1 × 10^6^ cells) were resuspended in 100 μl of supplemented solution of the Human Stem Cell Nucleofector Kit 1 (Lonza) containing a combination of the px458 plasmid targeting each gene and the ssODN and then electroporated using B-016 program of Nucleofector 2b (Lonza). The electroporated GSCs were cultured for 48 h followed by Single cell sorting of GFP-positive cells (SH800, SONY) into 96-well plates. For screening duplicated 96-well plates were lysed using QuickExtract DNA Extraction Solution (Epicenter) and the following primers were used to confirm the edited GSCs.

RELA – forward: 5’ – GGACATATGAGACCTTCCGC – 3’

RELA – reverse: 5’ – AGGGCTAGGTCAGTTCTCAG – 3’

### Immunofluorescence

GSC11 cells expressing wildtype or mutant RelA were seeded on coverslips to ∼70-80% confluency was allowed to attach for 12 hr followed by TNF-α (20ng/ml) treatment for 20 minutes and washed with PBS. Cells were then washed in cold PBS and fixed in 4% PFA for 15 min at RT followed by permeabilization with 0.3% Triton X-100 for 10 min at RT. Coverslips were then blocked by incubation in 2% BSA in PBS at RT for 30 min followed by 5% donor bovine serum (ThermoFisher Scientific) for 20 min. Blocked coverslips were then probed with antibodies detecting RelA (Cell Signaling Technology) in PBS containing 2% BSA and incubated in a humified chamber overnight at 4C. Next, fluorochrome-conjugated secondary antibody (Invitrogen, #A32731) working concentration in PBS containing 2% BSA was added to the coverslips and incubated for 2 hrs at RT in the dark. Coverslips were mounted with Fluoro-Gel II with DAPI (Electron Microscopy Sciences). Imaging was conducted using a Keyence microscope att 20X or 40X magnification.

### RNA-seq data analysis

RNA sequencing was performed by Novogene Corporation Inc, Sacramento, CA (read length: paired end 150). Analysis was performed as previously described (54). Fastq file quality was ensured with FastQC, and reads were aligned to the hg38 index using STAR. After alignment, the quality of the alignment was confirmed by assessing the final log output of STAR. Index and bigwig files were generated from the BAM files with samtools and bamCoverage, respectively. Each sample was uploaded to IGV to assess the quality and features of the reads mapping to the genome. The Subread package was used to generate count files, and these raw counts were used as input files for differential gene expression (DEG) analysis. Genes with less than 50 total counts across all samples were filtered out for DEG analysis. All above packages were downloaded and maintained with Conda package manager.

In R, DESeq2, pheatmap, and EnhancedVolcano packages were used for DEG analysis, hierarchical clustering visualized in heatmaps, and volcano plots, respectively. Volcano plotted genes by log-fold change and p-values. Initial DEGs were determined with a p_adj_ value of < 0.05 (BH correction). For Gene Ontology (GO) analysis, DEGs were further filtered to be decreased by at least 2-fold. GO analysis was done using Metascape (55). The gene list Venn diagram was generated using the online tool found at https://www.bioinformatics.org/gvenn/index.html. Lastly, the Gene Set Enrichment Analysis (GSEA) software was used to enrichment analysis for GBM subtypes (56–58). Input gene list files were generated with DESeq2 in R according to their GSEA documentation.

### Chromatin immunoprecipitation-qPCR

Chromatin immunoprecipitation was performed according to manufacturer instructions (Active motif) with the following modifications. Chromatin was sheared in diluted lysis buffer to 200 to 500 bp using a Covaris M220 Focused-Ultrasonicator with the following parameters: 3 minutes, peak incident power 75, duty factor 10%, 200 cycles/burst. Antibodies for ChIP were obtained from commercially available sources: anti-BRD2 (Cell Signaling Technology, #5848), anti-RelA/p65 (Cell Signaling Technology, #8242), anti-BRD4 (Cell Signaling Technology, #13440). Five percent of the chromatin was not exposed to antibody and was used as control (input). For ChIP-qPCR analysis, DNA quantity for each ChIP sample was normalized against input DNA.

### Quantitative real-time PCR

RNA was extracted with the RNeasy Plus kit (Qiagen, #74134) according to the manufacturer’s instructions. Reverse transcription of mRNA was performed using 3-5 μg RNA with RNA to cDNA EcoDry Premix (Takara, #639549). For real-time PCR analysis, 1 μL of cDNA (10 ng of starting RNA) was amplified per reaction using the iTaq Universal SYBR Green Supermix (Bio-Rad, #1725124) and the Bio-Rad CFX96 qPCR system.

### shRNA mediated knockdown

For lentivirus production, 293T cells were transfected with BRD2 shRNA (sigma), psPAX2 and pMD2.G packaging constructs using TransIT-VirusGEN (Mirus). Supernatants containing high titer lentiviruses were collected at 48 and 72 hr after transfection and were filtered through a 0.45 µm cellulose acetate filters before use. Viral preparations were then purified by LentiX concentrator (Takara Bio). GSCs were infected with lentivirus and after 24 hours of incubation at 37°C, the supernatant containing virus was replaced by fresh culture media. Infected cells were selected by puromycin.

### Matrigel invasion Assay

GSCs (1 x10^5^ cells) were suspended in serum-free culture medium and seeded into 24-well Transwell inserts (8.0 mm). Medium with indicated factors was added to the remaining receiver wells. After 16 - 24 h, the invaded GSCs were fixed and stained with crystal violet (0.05%, Sigma), and then counted as cells per field of view under microscope (Keyence BZ-X700).

### Intracranial injection

Animal research experiments were conducted under the regulations of the UCSD Animal Care Program, protocol number S00192M. A total of 1 × 10^5^ cells in a 5-μL volume was injected intracranially into 4- to 5-wk-old athymic nude mice using a stereotactic system. Tumors were allowed to establish for indicated weeks before any treatment, and engraftment of tumors was quantitatively confirmed via FMT signal intensity at the onset of neurological symptoms in the control groups. Tumor growth was monitored using the FMT 2500 fluorescence tomography system (PerkinElmer). For drug treatment studies, vehicle (DMSO) or 10 mg/kg GSK620 resuspended in vehicle was administered once every 2 days to mice via oral gavage. Mice were euthanized in accordance with our institutional guidelines for animal welfare and experimental conduct at University of California at San Diego.

### Immunohistochemistry

Formalin-fixed, paraffin-embedded (FFPE) tissue sections were prepared by the Histology Core Facility at UCSD pathology. Immunohistochemistry was performed according to standard procedures. Antigen was retrieved by boiling slides in 0.01 M of sodium citrate (pH 6.0) in a microwave for 15 min. Sections were incubated with primary antibody at 4°C overnight, followed by incubation with biotinylated secondary antibodies at room temperature for 30 min. Representative images from each immunostained section were taken with a Keyence BZ-X700 microscope and analyzed with BZ-X Analyzer Keyence software.

### Cell Growth Assay

5 replicates were plated in each well of black-walled, clear-bottom 96-well plates. Cell growth was analyzed using ATPlite 1step assay kit (PerkinElmer 6016731) following the manufacturer’s instructions.

### Pharmacokinetic studies

Whole blood from mice was centrifuged to isolate plasma. GSK620 was isolated by liquid-liquid extraction from plasma: 50 µL plasma was added to 2 µL internal standard and 3-fold volume acetonitrile. Mouse brain tissue was washed with 2 mL cold PBS and homogenized using a sonicator with fresh 2 mL cold PBS. GSK620 was then isolated and reconstituted in a similar manner by liquid-liquid extraction: 100 µL brain homogenate was added to 2 µL internal standard and 3-fold volume acetonitrile. After vortex mixing, the samples was centrifuged. The supernatant was removed and evaporated by a rotary evaporator and reconstituted in 100 µL 50:50:0.1 water:acetonitrile:formic acid.

### GSK620 detection

Chromatographic separations were performed on a 100 x 2.1 mm Phenomenex Kinetex C18 column (Kinetex) using the 1290 Infinity LC system (Agilent). The mobile phase was composed of solvent A: 0.1% formic acid in Milli-Q water, and B: 0.1% formic acid in acetonitrile. Analytes were eluted with a gradient of 5-95% B (1-15 min), 95% B (15-20 min), and then returned to 5% B for 5 min to re-equilibrate between injections. Injections of 20 µL into the chromatographic system were used with a solvent flow rate of 0.10 mL/min.

Mass spectrometry was performed on the 6460 triple quadrupole LC/MS system (Agilent). Ionization was achieved by using electrospray in the positive mode and data acquisition was made in multiple reactions monitoring (MRM) mode. Two MRM transitions were used for GSK620: m/z 325→ 169 and 325→ 247 with fragmentor voltage of 85V, and collision energy of 17 and 5 eV, respectively. Analyte signal was normalized to the internal standard and concentrations were determined by comparison to the calibration curve (0.5, 5, 50, 250, 500, 2000 nM). GSK620 brain concentrations were adjusted by 1.4% of the mouse brain weight for the residual blood in the brain vasculature as described by Dai et al.(59).

### Statistical Analyses

Statistical analyses were performed using GraphPad Prism 9 software. Data sets were analyzed by unpaired t-test or multiple comparisons by one-way ANOVA or two-way ANOVA according to the experiment. In figures, a single asterisk indicates *P* < 0.05, double asterisks indicate *P* < 0.001, and triple asterisks indicate *P* < 0.0001.

## Acknowledgments

We are grateful to Dr. Andrew Shiau for GSK620. We thank Dr. Krishna Bhat for GBM-PDX neurospheres, Dr. Antonio Iavarone and Dr. Simona Migliozzi for assistance in bio-informatic analysis, Dr. Donald Pizzo for IHC sections. This work was supported by R01NS080939, R56NS080939, and R01CA258248 to F.B.F., the Japan Society for the Promotion of Science Overseas Research Fellowship to S.M., and the Kurland Family Foundation and P50CA211015 to D.A.N

## Author contributions

R.V and F.F. conceived and designed the study. R.V., S.M., B.T., D.K., B.M.J., N.N. and P.P. conducted the experiments. R.V analyzed and interpreted the data. F.F. supervised all aspects of the study. R.V. and F.F. wrote the manuscript with input from other authors.

## Data availability

All data will be available upon request.

## Notes

### Competing Interest Statement

The authors have declared no competing interest.

