## Supplemental Material for "Glioblastoma Mesenchymal Transition and Invasion are Dependent on a NF-κB/BRD2 Chromatin Complex"

### Supplemental Figure 1

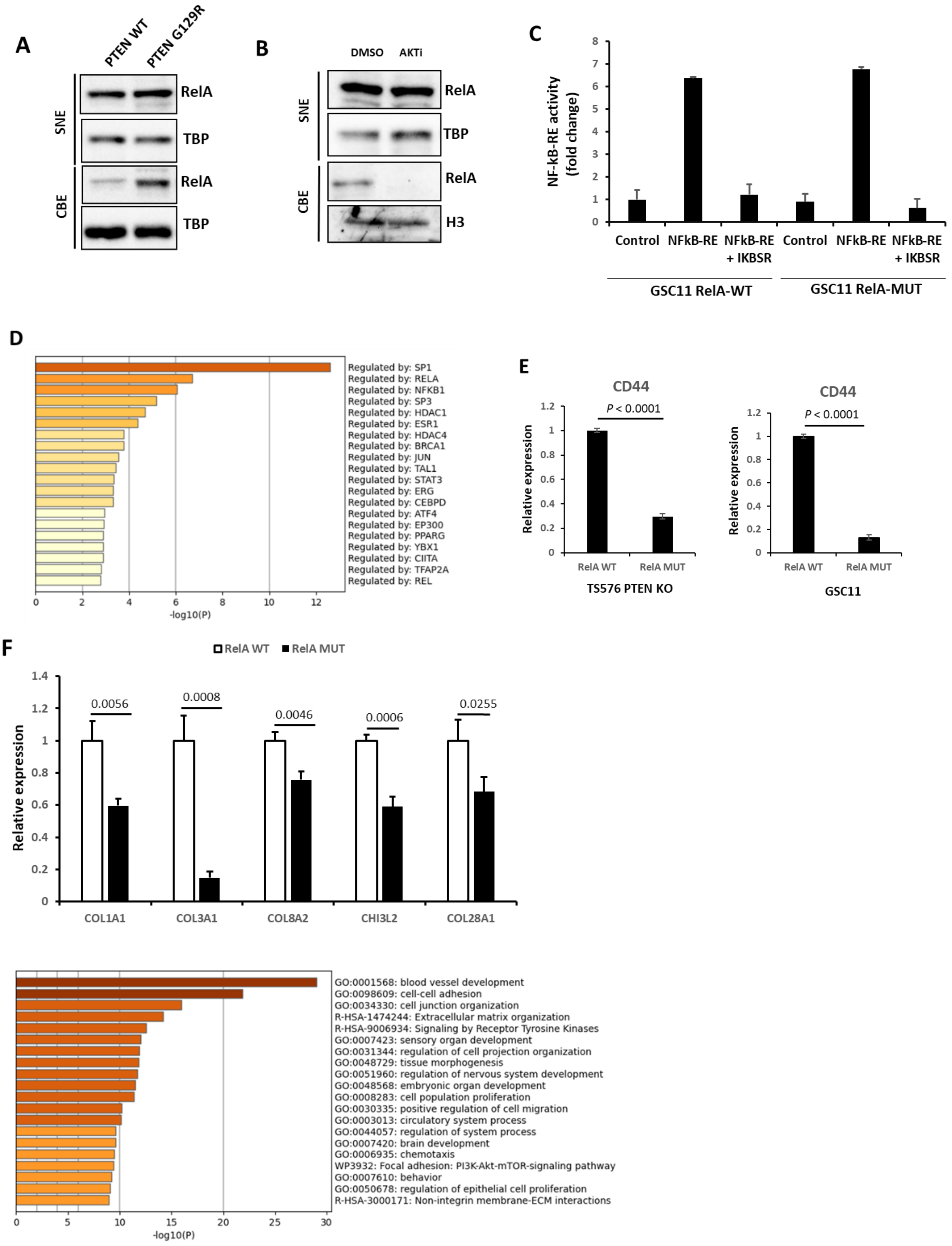

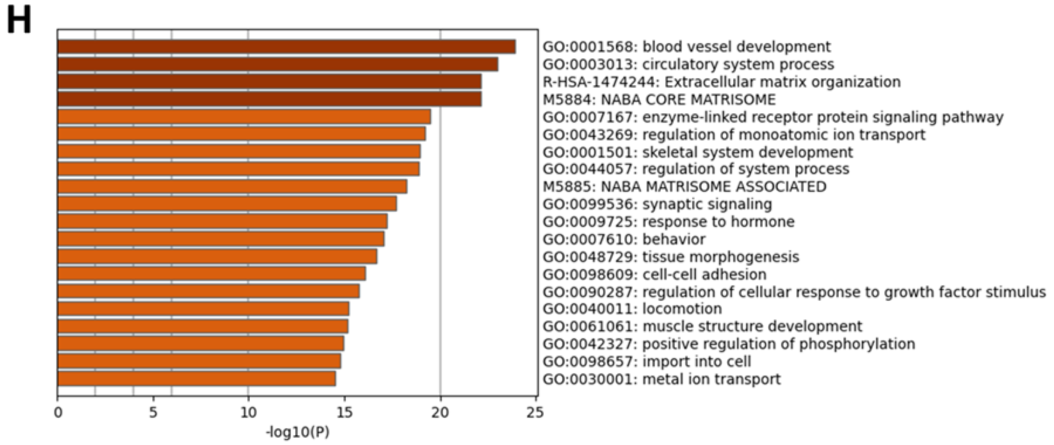


**Figure S1**. RelA K310 acetylation recruits BRD2 and BRD4 to chromatin and regulates ECM gene expression in PTEN deficient GBM. **A)** Western blot analysis of RelA in SNE and CBE fractions of U87 cells stably expressing wildtype or phosphatase dead (G129R) PTEN. **B)** Western blot analysis of RelA in SNE and CBE fractions of GSC11 cells treated with DMSO or Ipatasertib (AKTi, 1uM) for 24h. TBP and H3 were used as loading controls for SNE and CBE fractions respectively. **C)** Luciferase assay for NF-κB activity in RelA wildtype or mutant GSC11 cells with or without IKBSR expression**. D)** Regulatory transcription factors enriched in RelA Mutant GSC11 downregulated genes. **E)** qPCR analysis of CD44 expression in RelA WT and RelA MUT cells in GSC11 and TS576 PTEN KO cells. Data represents mean ± SD. **F)** qPCR analysis of ECM gene expression in RelA WT and RelA MUT GSC11 cells. Data represents mean ± SD. **G)** GO analysis of downregulated genes in TS576 PTEN KO/ RelA-MUT cells compared to wildtype TS576 PTEN KO cells. **H)** GO analysis of downregulated genes in TS576 PTEN KO cells treated with CBP/P300 inhibitor (A-485) for 24h.

**Supplemental Figure 2**

**A**

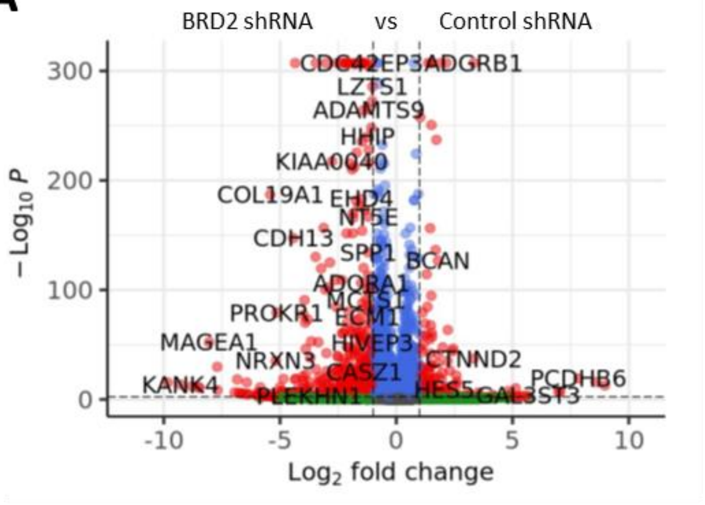


**Figure S2**. **A)** Volcano plot showing the differential gene expression profiles of GSC11 BRD2 shRNA vs GSC11 control shRNA expressing cells. Fold change was plotted as log2(fold change) for each gene relative to its false discovery rate (−log2[FDR]).

### Supplemental Figure 3

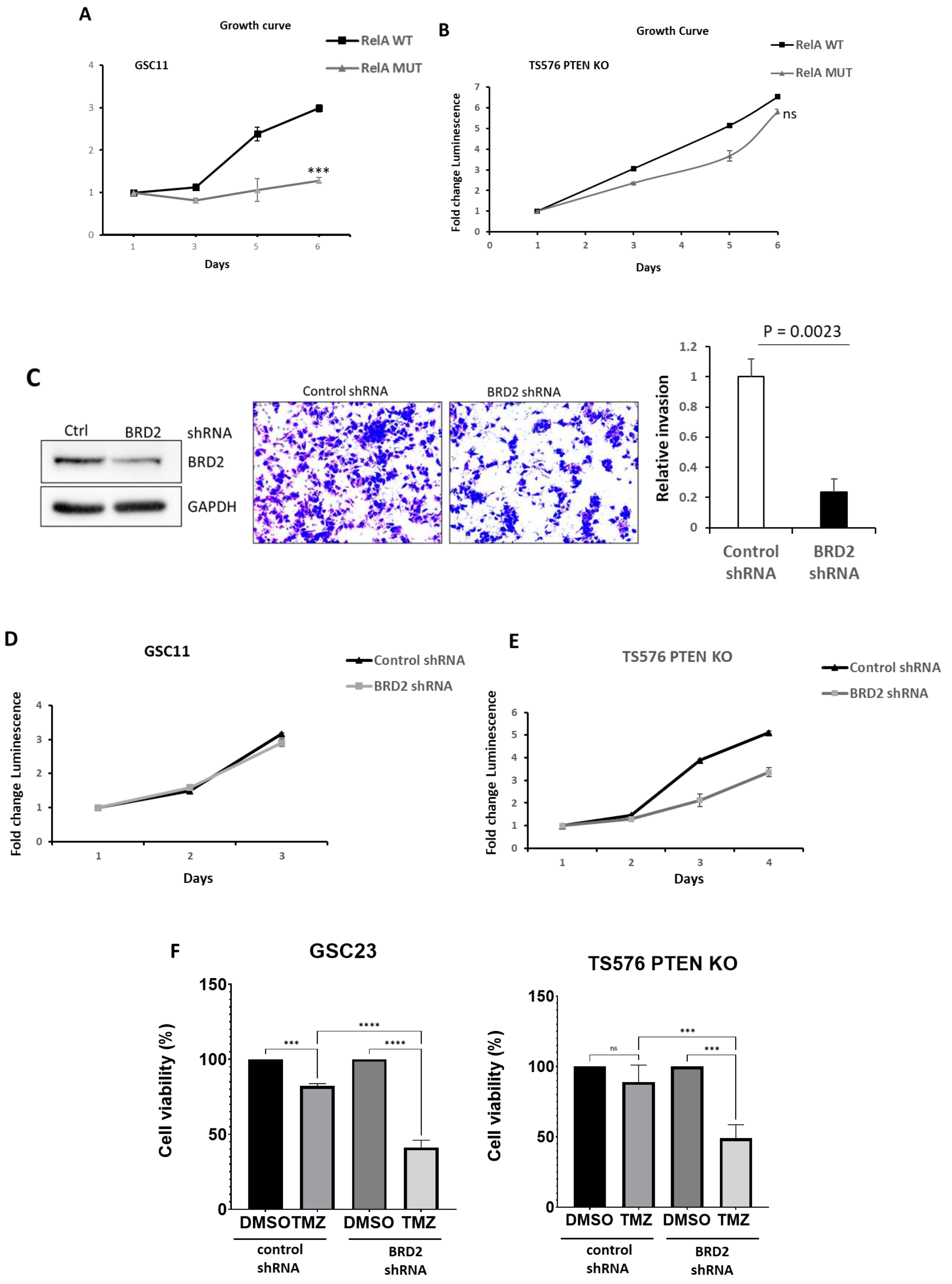

**Figure S3**. Cell growth and invasion analysis of wildtype and mutant RelA cell lines. ATPlite assay showing cell growth differences in **A)** RelA WT and RelA MUT GSC11 cells and in **B)** RelA WT and RelA MUT TS576 PTEN KO cells. **C)** Immunoblot analysis and matrigel invasion assay of TS576 PTEN KO expressing control or BRD2 shRNAs. Data represents mean ± SD. **D-E)** ATPlite assay showing cell growth differences in control and BRD2 shRNAs expressing GSC11 and TS576 PTEN KO cells. **F)** Control or BRD2 shRNAs expressing GSCs were treated with 100uM TMZ for 24 h and cell viability was assessed by ATPlite assay. Data represents mean ± SD

### Supplemental Figure 4

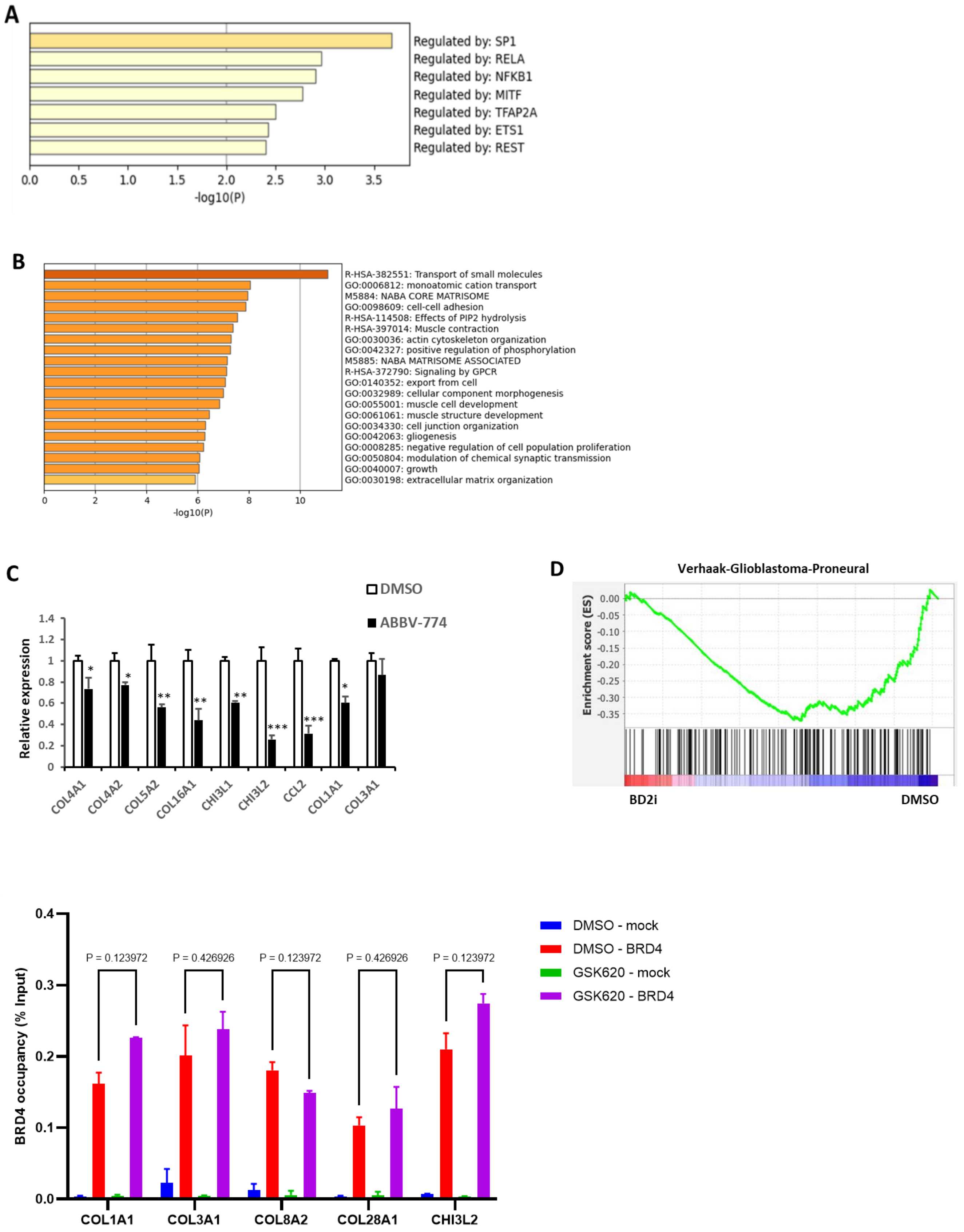

F

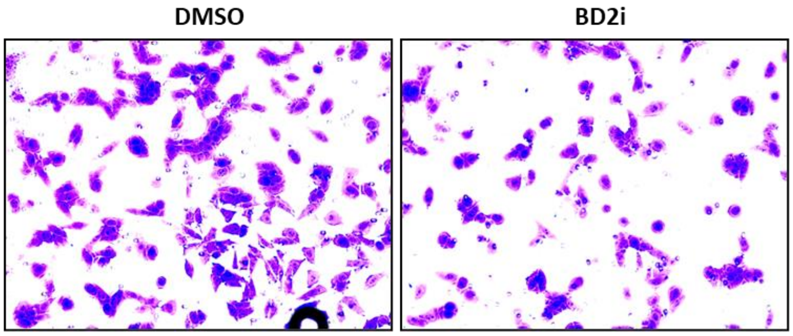


**Figure S4**. BET-BD2 inhibitors mimic RelA K310R mutation in regulating ECM gene expression and invasion. **A)** Regulatory transcription factors enriched in GSK620 treated GSC11 downregulated genes. **B)** GO analysis of downregulated genes in TS576 PTEN KO cells treated with GSK620 (0.5uM for 24h) compared to DMSO treated cells. **C)** qPCR analysis of ECM genes in ABBV-744 treated GSC11 cells. Data represents mean ± SD. **D**) GSEA enrichment enrichment plots of DMSO or BD2i gene lists versus PN gene expression is shown. **E)** ChIP-qPCR with anti-BRD4 on the promoters of ECM genes in DMSO and BD2i treated GSC11 cells. For ChIP assays, bar graphs indicate fold enrichment of BRD2 over the input (n=3 biological samples with three replicates each). **F)** Matrigel cell invasion of GSC267 cells treated with DMSO or BD2i for 24h were examined by transwell-chamber assays.

**Supplemental Figure 5**

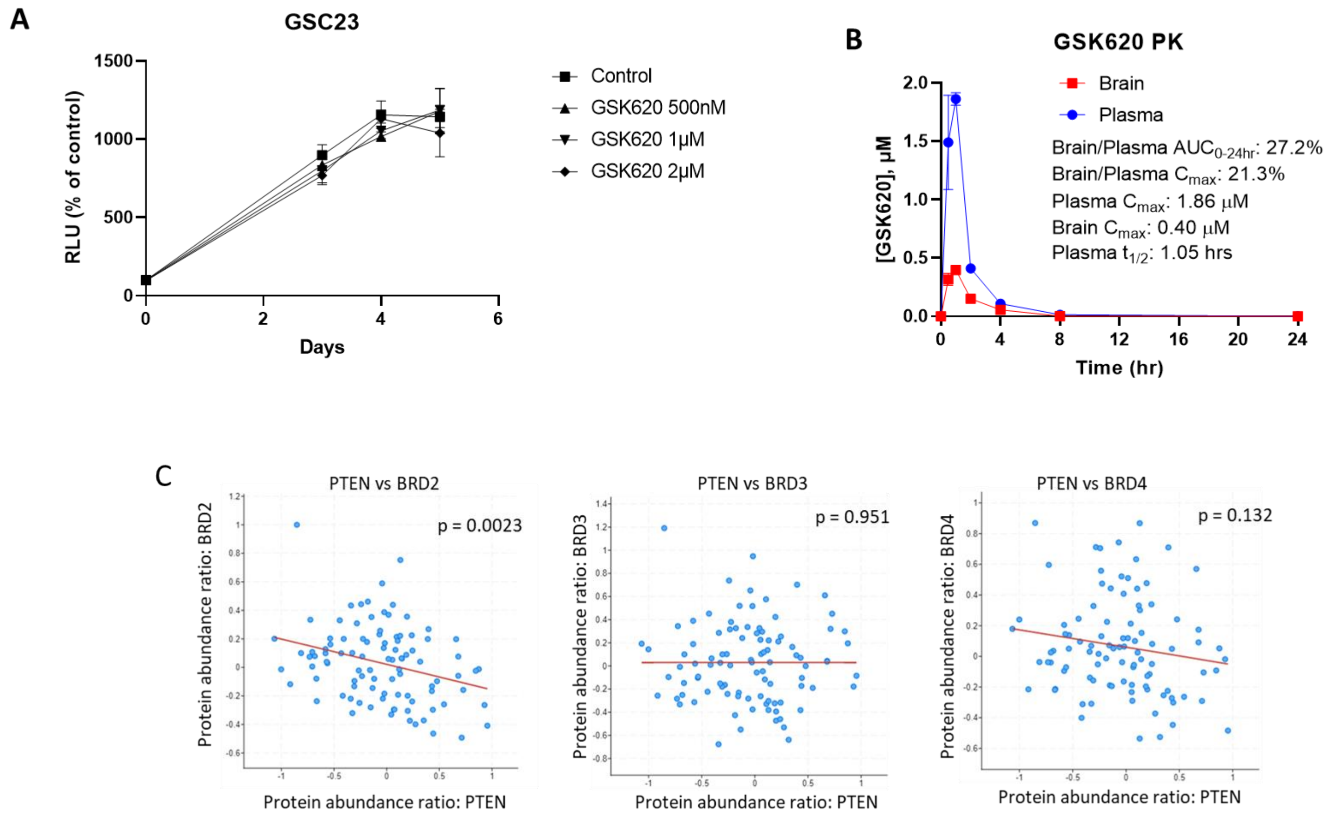


**Figure S5**. **A)** Effect of GSK620 on GSC23 *in vitro* cell growth by ATPlite assay. **B)** PK analysis of GSK620 in nude mice. **C)** Pearson correlation analysis of BRD2, BRD3 and BRD4 protein levels with PTEN in GBM patient samples (cBioportal). n=99.
